# Targeting CXADR-mediated AKT signaling suppresses tumorigenesis and enhances chemotherapy efficacy in Ewing sarcoma

**DOI:** 10.1101/2025.08.07.669049

**Authors:** Alina Ritter, Malenka Zimmermann, Florian H. Geyer, Pablo Táboas, David Obermeier, Martha J. Carreño Gonzalez, Kimberley M. Hanssen, Tobias Faehling, Jana Siebenlist, Annika Jeschke, Laura Romero-Pérez, Roland Imle, Ana Banito, Enrique de Álava, Wolfgang Hartmann, Uta Dirksen, Thomas G. P. Grünewald, Florencia Cidre-Aranaz

**Affiliations:** Hopp Children’s Cancer Center Heidelberg (KiTZ), Heidelberg, Germany; National Center for Tumor Diseases (NCT), NCT Heidelberg, a partnership between DKFZ and Heidelberg University Hospital, Heidelberg, Germany; Division of Translational Pediatric Sarcoma Research, German Cancer Research Center (DKFZ), German Cancer Consortium (DKTK), Heidelberg, Germany; Faculty of Medicine, Heidelberg University, Heidelberg, Germany; Department of Internal Medicine V, Heidelberg University Hospital, Heidelberg, Germany; Department of Pediatric Oncology and Hematology, Hospital Sant Joan de Déu, Barcelona 08950, Spain; Instituto de Biomedicina de Sevilla, IBiS/Hospital Universitario Virgen del Rocio/CSIC/Universidad de Sevilla/CIBERONC; Department of Human Anatomy and Embryology, Faculty of Medicine, University of Seville, Seville, Spain; Soft-Tissue Sarcoma Junior Research Group, DKFZ, Heidelberg, Germany; Department of Pediatric Oncology, Hematology and Immunology, Heidelberg University Hospital, Heidelberg, Germany; Department of Normal and Pathological Histology and Cytology, Faculty of Medicine, University of Seville, Seville, Spain; Department of Pathology, University Hospital Virgen del Rocío, Pathology Unit, Seville, Spain; Gerhard-Domagk-Institute of Pathology, University of Muenster, Muenster, Germany; Pediatrics III, AYA Unit, West German Cancer Centre, University Hospital Essen, Essen, Germany; German Cancer Consortium (DKTK), partner site Essen, Essen, Germany; Institute of Pathology, Heidelberg University Hospital, Heidelberg, Germany

## Abstract

Distant metastasis is the leading cause of mortality in Ewing sarcoma (EwS) – a malignant bone or soft-tissue cancer mainly affecting children, adolescents, and young adults. Despite continuous efforts in understanding its pathogenesis, the molecular mechanisms driving EwS metastasis remain poorly understood, thus limiting the potential for therapeutic progress. Here, we identify the tight junction component Coxsackievirus and Adenovirus receptor (CXADR) as a critical regulator of cancer progression and metastasis in EwS. Differential gene expression analysis of patient tumors from two independent cohorts revealed that elevated *CXADR* levels are associated with metastatic disease and poor overall survival. In functional experiments, conditional CXADR knockdown reduced the growth of EwS cell line models in vitro, and suppressed local tumorigenesis. Notably, CXADR knockdown completely abrogated metastasis formation in vivo. Integration of transcriptome profiling and mechanistic studies uncovered that CXADR promotes the activation of AKT signaling, likely through complex formation with PTEN. Consequently, pharmacological targeting of AKT using the FDA-approved pan-AKT inhibitor Capivasertib showed CXADR-dependent cytotoxicity, with enhanced efficacy if combined with the EwS standard-of-care chemotherapeutic agent Vincristine.

Collectively, our findings establish CXADR as a prognostic and predictive biomarker in EwS, highlighting AKT inhibition combined with chemotherapy as a promising strategy for patients with high CXADR expression. Together, these findings support a precision medicine approach combining molecular stratification and targeted therapies to improve patient outcomes in metastatic EwS.

## INTRODUCTION

Metastases are responsible for approximately two-thirds of all cancer-related deaths in solid tumors (Dillekås et al., 2019). In Ewing sarcoma (EwS) – the second most common malignant bone or soft-tissue cancer in children, adolescents, and young adults – the survival rate drops from 70–80% for localized disease to less than 30% for metastatic disease (Casali et al., 2018; Grünewald et al., 2018; Leavey et al., 2021). To date, the underlying mechanisms of metastasis in EwS have only been partially understood (Bull et al., 2025; Grünewald et al., 2018), which may constitute one of the main reasons for the lack of improvement in patient outcomes (Setty et al., 2023). Hence, the identification of genes that influence the progression of EwS is of central importance for a better understanding of the disease and for future development of novel biomarkers and targeted therapies.

In carcinomas, the metastatic process is mediated by epithelial-to-mesenchymal transition (EMT), during which cells reduce their proliferative capacity and acquire increased migratory and invasive potential (Kalluri and Weinberg, 2009). In sarcomas, it has been proposed that tumor cells may reside in a metastable state where they could easily switch towards a more mesenchymal or epithelial accentuation depending on internal/external cues (Cidre-Aranaz et al., 2022; Franzetti et al., 2017; Sannino et al., 2017). These signals often rely on activation of critical pathways, which in sarcomas include regulation of WNT, MEK-ERK, and AKT, among others (Cano et al., 2000; Hawkins et al., 2020; Pedersen et al., 2016; Sannino et al., 2017). However, the precise mechanisms and their central effectors in EwS remain poorly defined. Here, we propose Coxsackievirus and Adenovirus receptor (CXADR) – a viral receptor and component of the tight junction complex (Nilchian et al., 2019; Zhang and Lui, 2023) – as a variably expressed activator of AKT signaling in EwS. CXADR consists of an extracellular domain responsible for cell-cell adhesion, a transmembrane domain, and an intracellular domain, which facilitates interactions with other signaling proteins like ZO-1, MAGI-1, or LNX via its PDZ-binding site (Matthäus et al., 2017; Nilchian et al., 2019; Ortiz-Zapater et al., 2017; Zhang and Lui, 2023).

In this study, we report that in EwS, CXADR inhibition results in a strong reduction of tumorigenic features and an abrogation of metastatic phenotypes in vitro and in vivo due to reduced AKT signaling. In EwS patients, higher expression levels of CXADR are associated with worse survival and higher metastatic potential. Indeed, in CXADR-high conditions, we show a synergistic pharmacological inhibition of AKT combined with standard-of-care chemotherapy, which poses an opportunity for the application of targeted therapeutics based on specific patient stratification according to their CXADR expression.

## RESULTS

### Identification of CXADR as a potential key regulator of metastasis in EwS

To identify clinically relevant candidate genes potentially promoting metastasis in EwS, we performed a differential gene expression (DGE) analysis of tumors derived from molecularly confirmed EwS patients displaying extreme phenotypes, i.e. worst outcome (WO, presence of metastatic spread, and death-of-disease during the follow-up period) in direct comparison with tumors derived from patients displaying best outcome (BO, as defined by localized disease at diagnosis, and long-term survival during the follow-up period with no evidence of disease). To prevent potential confounding effects due to prior therapy, all samples selected belonged to patients who had seen no previous treatment. Thus, we devised a pipeline where RNA-seq data of a first cohort comprising these 12 tumor samples (n_WO_ = 7; n_BO_ = 5) were compared using the DESeq2 algorithm in R (**Fig. 1a**). DGE analysis yielded a set of 369 differentially expressed genes (DEGs) (**Fig. 1a**). Since, from a translational and pharmacological perspective, it is generally more feasible to target proteins that are highly expressed than to attempt re-expression of silenced ones, we further filtered these 369 DEGs to retain only those upregulated in metastatic disease. Additionally, the data of a second, independent cohort of 166 EwS patients was leveraged to filter for those genes being associated with worse patient overall survival (**Fig. 1a**). Next, the top 15 candidates were subjected to gene ontology (GO) enrichment analysis for GO terms associated with metastasis (**Supp. Table 1**), which resulted in three genes, namely *ANKLE1*, *BRSK2*, and *CXADR* (**Fig. 1a**). Lastly, we prioritized the candidates according to their expression in EwS. For this, the expression of each candidate gene was assessed in 18 EwS cell lines included in the Ewing Sarcoma Cell Line Atlas (ESCLA) project (Orth et al., 2022) for the highest average expression. This analysis revealed the Coxsackievirus and Adenovirus receptor (*CXADR*) as being highly expressed in EwS cell lines and in patients presenting with metastatic disease. Its expression was associated with metastatic-related GO terms, and correlated with poorer patient overall survival (**Fig. 1b,c**).

**Figure 1.**
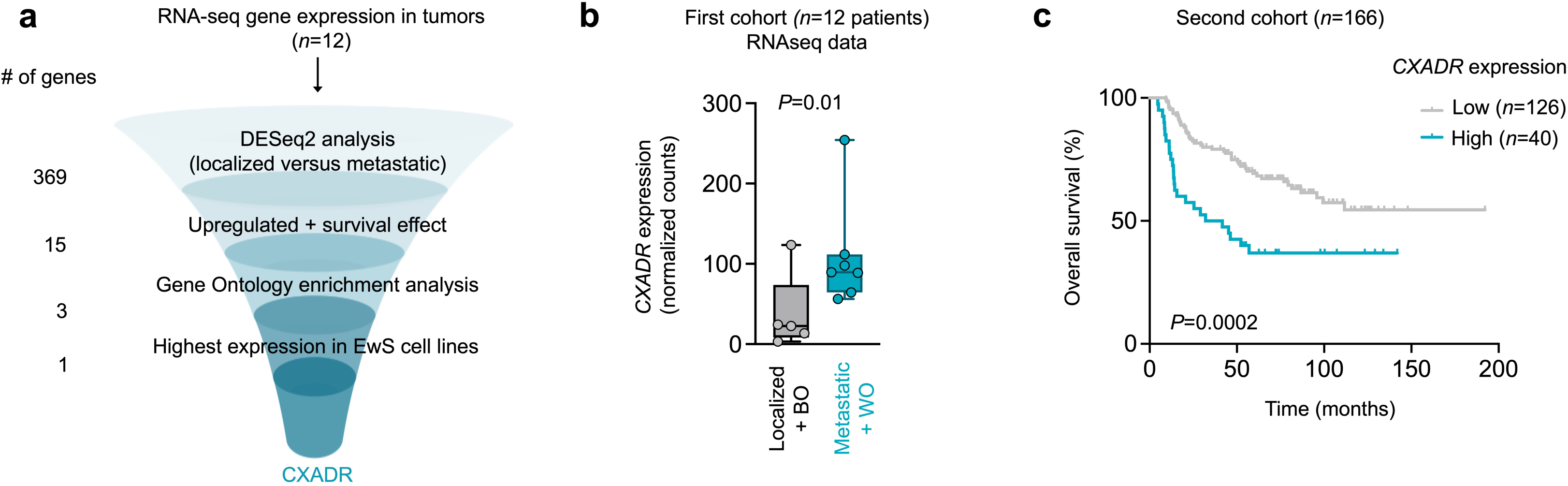
Identification of *CXADR* as a potential key mediator for metastasis in EwS. **(a)** Systematic pipeline identifies *CXADR* as a prognostically-relevant, metastasis-related, upregulated gene in EwS. **(b)** Comparison of the normalized counts of *CXADR* (RNA-seq) in best outcome (BO, localized disease and survival in the observation period with no evidence of disease) versus worst outcome (WO, metastatic disease and death in the observation period) in the first patient cohort. *P*-value was determined using DESeq2 analysis. **(c)** Overall survival in a second cohort of 166 EwS patients according to *CXADR* expression. Best percentile was determined as cutoff between groups. Log-rank (Mantel-Cox) test.

To exclude the possibility that the patients with metastatic status, who generally presented higher expression of *CXADR* (first cohort, **Fig. 1b**), were driving the survival effect observed in the second cohort (**Fig. 1c**), we employed extended clinical metadata from the second cohort to assess the *CXADR* expression of EwS patients according to their sample origin (primary or metastatic lesions). As expected, *CXADR* was significantly higher expressed in metastasis-derived samples (*P* = 0.0427; **Supp. Fig. 1a**). Interestingly, the exclusion of these samples from the survival analysis still revealed a strong and significant association of high *CXADR* expression with worse patient overall survival (*P* = 0.0001; **Supp. Fig. 1b**). Similarly, even after exclusion of patients with metastatic disease from the analysis, the significant association of high *CXADR* expression with worse overall survival was preserved (*P* = 0.02; **Supp. Fig. 1c**), suggesting that high *CXADR* is already a feature of the primary tumors that will result in a worse patient outcome.

Collectively, these results suggest that *CXADR* is highly expressed in metastasis-derived samples in EwS and constitutes a risk factor for a worse overall survival in EwS patients, even in localized disease.

### CXADR promotes tumorigenesis in EwS

To explore the role of *CXADR* in EwS, we first inhibited its expression in two EwS cell lines selected because of their relatively high *CXADR* expression (MHH-ES-1, SK-N-MC). A strong siPOOL-mediated *CXADR* knockdown (KD) for 72 h resulted in a significant reduction of cell proliferation in both cell lines (*P*_MHH-ES-1_ = 0.0286; *P*_SK-N-MC_ = 0.0022; **Supp. Fig. 2a,b**). To validate these results with an alternative RNA interference technology, we transduced MHH-ES1 and SK-N-MC EwS cell lines with doxycycline (DOX)-inducible shRNAs targeting either the 3’UTR region or the coding sequence (CDS) of *CXADR* or an untargeted shRNA control sequence (**Supp. Table 2**). KD efficacy was confirmed in all cell lines upon addition of DOX (1 µg/mL) for 72 h at mRNA and protein levels and was not observed in the control conditions (**Fig. 2a,b, Supp. Fig. 2c**). In agreement with our previous results, conditional *CXADR* KD resulted in a significant inhibition of cell proliferation (**Fig. 2c**, *P*_MHH-ES-1_ = 0.0079; *P*_SK-N-MC_ = 0.0079), which was not observed in controls. Concordantly, cell cycle analysis of both cell lines upon *CXADR* KD showed an arrest in the G1 phase (**Supp. Fig. 2d**). Similarly, *CXADR* inhibition drastically impaired 2D clonogenic growth (**Fig. 2d**, *P*_MHH-ES-1_ = 0.0079; *P*_SK-N-MC_ = 0.0079) and 3D-spheroidal growth (**Fig. 2e**, *P*_MHH-ES-1_ = 0.0286; *P*_SK-N-MC_ = 0.0286) in both EwS cell lines. To further test this phenotype in vivo, we assessed local tumor growth by injecting MHH-ES-1 and SK-N-MC cells carrying a DOX-inducible KD of *CXADR* using two different shRNAs in the right flank of NSG mice. DOX-induced *CXADR* KD led to a strong and significant inhibition of tumor growth (*P*_MHH-ES-1_ = 0.000622; *P*_SK-N-MC_ = 0.0011; **Fig. 3a**). Concordantly, ex vivo analysis of resected tumor tissue revealed a significant reduction of tumor weight (*P*_MHH-ES-1_ = 0.0019; *P*_SK-N-MC_ = 0.0005; **Fig. 3b**), which was accompanied by a significant reduction of the mitotic index (*P*_MHH-ES-1_ = 0.0079; *P*_SK-N-MC_ = 0.0079; **Fig. 3c**), an increase of the necrotic area (**Supp. Fig. 3a**) and of the overall number of apoptotic cells as quantified by cleaved caspase 3 (**Fig. 3d**) in immunohistological stainings.

**Figure 2.**
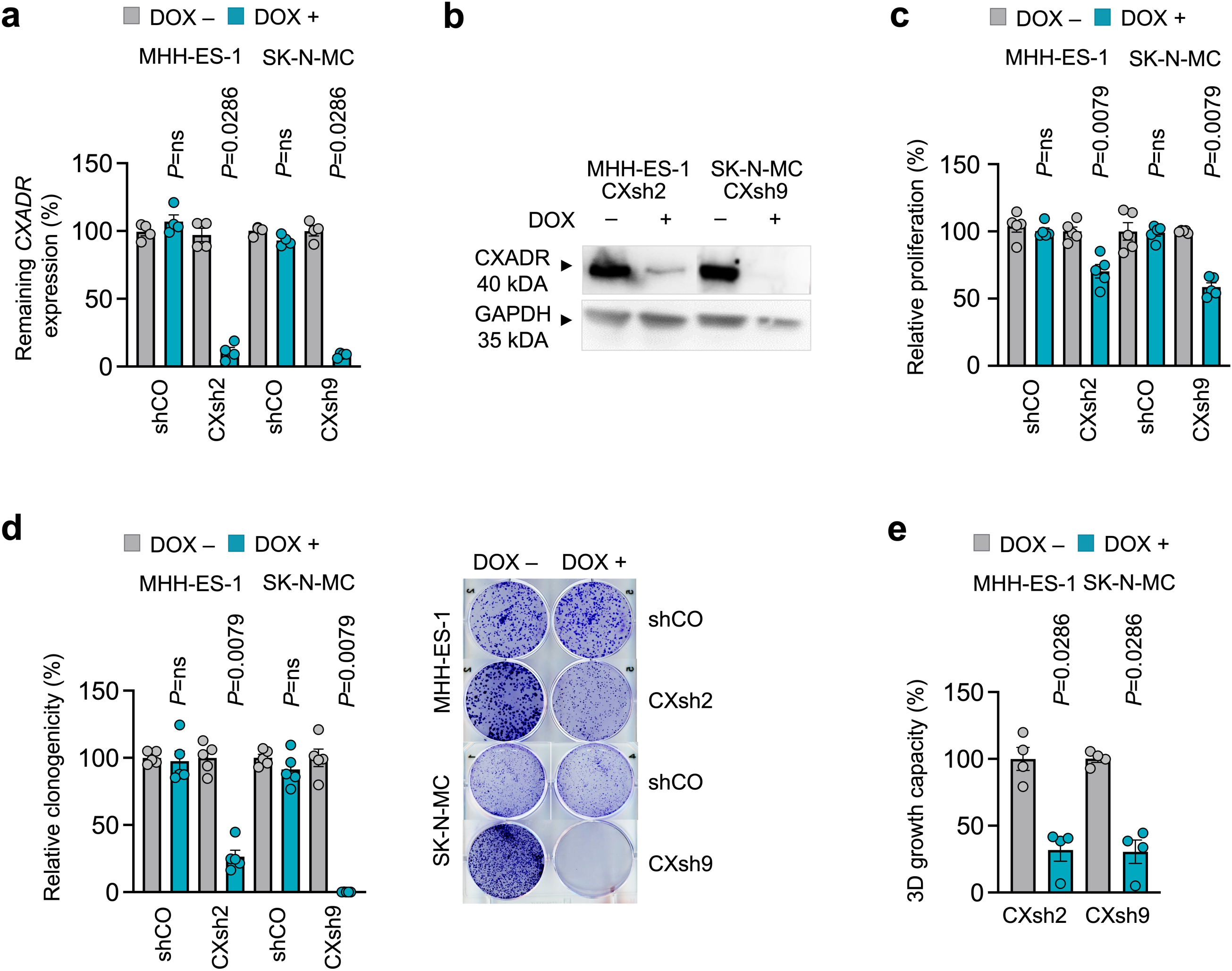
CXADR drives tumorigenesis in EwS in vitro. **(a)** Remaining mRNA expression of *CXADR* after 72 h of DOX treatment using two different shRNAs (sh2, c20; and sh9, c2) or non-targeting control (shCO) in two EwS cell lines. n = 4 independent biological replicates. Error bars represent SEM. Two-sided Mann-Whitney test. **(b)** Western blot showing *CXADR* KD at the protein level in two EwS cell lines after 72 h of DOX treatment. GAPDH was used as a loading control. **(c)** Proliferation assay after 72 h of DOX treatment. *n* = 5 independent biological replicates. Error bars represent SEM. *P*-values were determined using a two-sided Mann-Whitney test. **(d)** Left: quantification of clonogenicity of EwS cells upon KD of *CXADR* for 10–14 d. *n* = 5 independent biological replicates. Error bars represent SEM. *P*-values were determined using a two-sided Mann-Whitney test. Right: representative pictures. **(e)** Left: quantification of relative sphere formation capacity in two EwS cell lines upon *CXADR* KD. *n* = 4 independent biological replicates. *P-*values were determined using a two-sided Mann-Whitney test.

**Figure 3.**
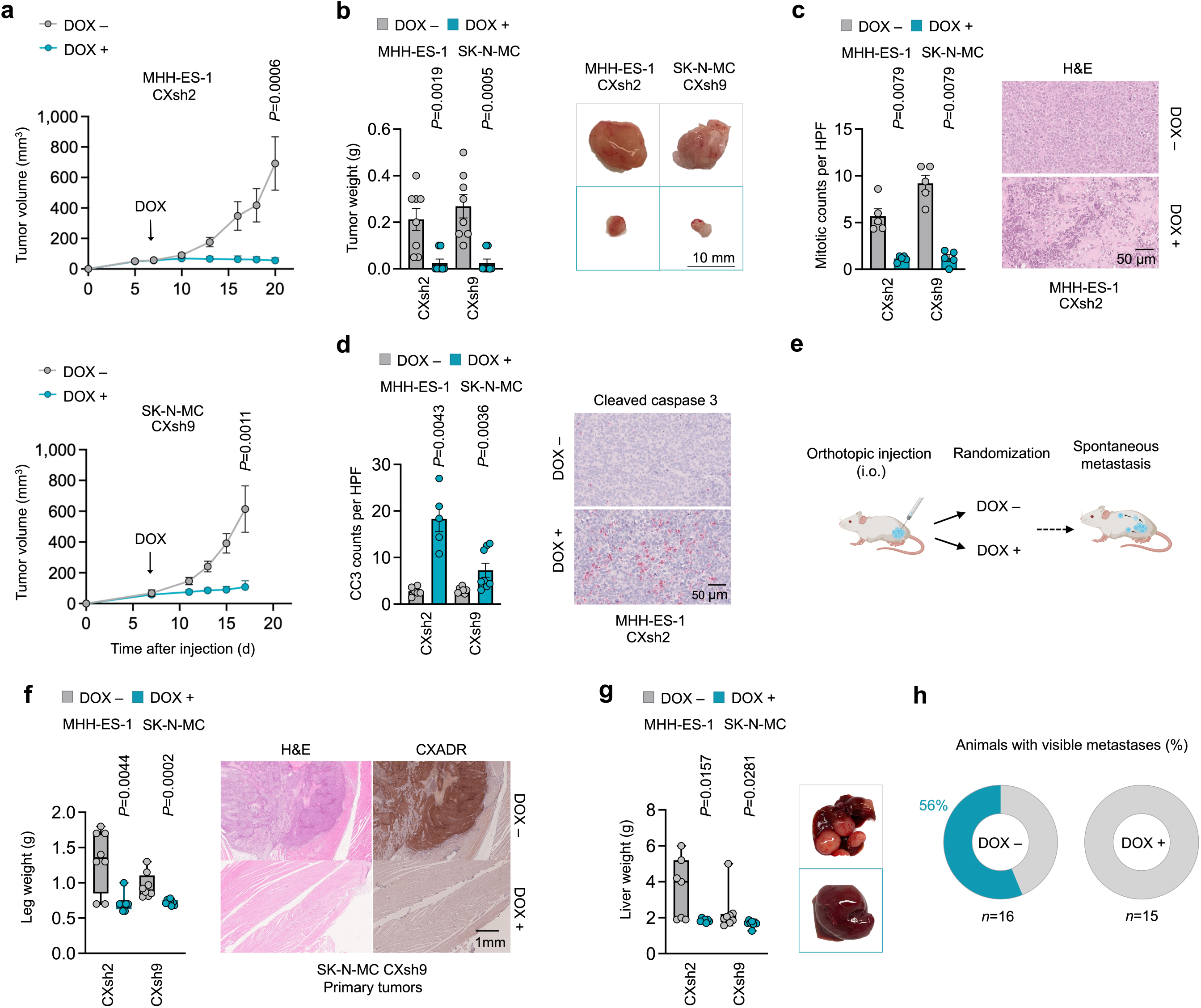
CXADR drives primary tumor growth and metastatic spread in EwS in vivo. **(a)** Growth curves of EwS subcutaneous xenografts of MHH-ES-1 and SK-N-MC cells containing a DOX-inducible KD for *CXADR* (arrow indicates start of DOX-treatment). Data are represented as means (*n* ≥ 7 animals/group). Error bars represent SEM. Two-sided Mann-Whitney test. **(b)** Left: weight of extracted primary tumor tissue. Error bars represent SEM. *P*-values were determined using a two-sided Mann-Whitney test. Right: representative images of extracted tumors for each experiment. **(c)** Mitotic counts per high power field (HPF) and representative HE pictures in two different EwS cell lines with two different shRNA constructs. *P*-values were determined using a two-sided Mann-Whitney test. **(d)** Cleaved-caspase 3 counts per HPF and representative pictures of two different EwS cell lines with two different shRNA constructs. *P*-values were determined using a two-sided Mann-Whitney test. **(e)** Schematic representation of experimental design: MHH-ES-1 or SK-N-MC EwS cell lines were orthotopically injected in the tibia plateau of NSG mice and treated with either DOX (DOX +) or sucrose (DOX–, control) to test for spontaneous metastatic spread to inner organs. **(f)** Left: comparison of ex vivo leg weight upon KD of *CXADR* in orthotopic mouse experiments. *P*-values were determined using a two-sided Mann-Whitney test. Right: Representative H&E histological images and immunohistological images stainings for CXADR of primary tumor growth in the legs of orthotopically xenografted mice. **(g)** Comparison of liver weight following *CXADR* KD in orthotopic experiments. *P*-values were determined using a two-sided Mann-Whitney-test. **(h)** End-point analysis of the presence of spontaneous metastases in inner organs. Pie charts depict the percentage of metastasis-free (grey) or -bearing (blue) animals in each condition, *n* represents the total number of animals.

Since *CXADR* is highly expressed in patients presenting with metastasis, we explored its contribution to the metastatic process in vivo using an orthotopic mouse model of spontaneous metastasis formation (Cidre-Aranaz et al., 2022). To that end, MHH-ES-1 and SK-N-MC EwS cells harbouring the DOX-inducible *CXADR* KD were injected in the right tibia of NSG mice that were subsequently subjected to either sucrose (control)- or DOX-treatment in the drinking water (**Fig. 3e**). Remarkably, in agreement with our previous results on local tumor growth (**Fig. 3a**), *CXADR* KD completely inhibited tumor growth at the primary location (leg) (**Fig. 3f**), which resulted in a prolonged survival of all DOX-treated animals (*P* = 0.0004; **Supp. Fig. 3b**).

In addition, *CXADR* KD resulted in a complete inhibition of metastatic spread to distant organs, as depicted by quantification of liver weight (*P*_MHH-ES-1_ = 0.0157; *P*_SK-N-MC_ = 0.0281; **Fig. 3g**) and percentage of animals with visible macrometastases (**Fig. 3h**).

In summary, *CXADR* silencing in EwS decreases proliferation, clonogenicity, and sphere formation, inhibiting local tumor growth and metastasis formation in vivo.

### CXADR mediates EMT-phenotypes via the AKT signalling pathway

To obtain insights into how CXADR mediates its phenotype in EwS, MHH-ES-1 and SK-N-MC EwS cells harboring a shRNA-mediated DOX-inducible *CXADR* KD were subjected to RNA-seq analyses and subsequent fGSEA using the Hallmark gene set at MSigDB (https://www.gsea-msigdb.org/gsea/msigdb/collections.jsp). Here, ^‘^epithelial-mesenchymal-transition’ emerged as one of the top negatively enriched signatures (NES = –1.65, *P_adj_* = 0.00335; **Fig. 4a; Supp. Fig. 4a**), suggesting that high CXADR expression promotes EMT. Further analysis of our RNA-seq data employing the highly curated c2 gene set (https://www.gsea-msigdb.org/gsea/msigdb/human/genesets.jsp?collection=C2) showed an overlap of five significantly regulated gene sets shared between the two EwS cell lines upon *CXADR* KD. These gene sets prominently included ‘PI3K-AKT-signaling-pathway’ (**Fig. 4b, Supp. Fig. 4b**), suggesting that CXADR may promote EMT via activation of the AKT pathway. To test this hypothesis, we performed time-course experiments using both EwS cell lines with a DOX-inducible shRNA against *CXADR*. Excitingly, western blot analyses showed that CXADR silencing led to a strong inactivation of AKT, as indicated by a decrease in pAKT protein levels (**Fig. 4c**). These observations were corroborated by a concomitant inactivation of AKT downstream targets such as GSK3ß (**Fig. 4c**). Although little is known about the precise mechanism of action of CXADR as a regulator of AKT, a recent report showed that CXADR may modulate AKT signalling by forming an inhibitory complex with PTEN and PHLPP2 in breast cancer (Nilchian et al., 2019). Thus, we inquired whether this regulatory mechanism is active in EwS. To address this question, we co-immunoprecipitated CXADR in MHH-ES-1 and SK-N-MC EwS cell lines. Western blot analysis using a PTEN-specific antibody confirmed that CXADR forms a complex with PTEN, implying that AKT signalling regulation through a CXADR-PTEN complex is also in effect in EwS (**Fig. 4d**). To probe whether this potential CXADR-regulation of the AKT-pathway may also be valid in EwS patients, we generated a custom-made signature consisting of five genes associated with downregulated AKT signaling, including *PTEN* (‘AKT_down’, **Supp. Table 3**), and interrogated our second EwS patient cohort (n = 166) using single-sample gene-set enrichment analysis (ssGSEA). Strikingly, *CXADR* expression significantly correlates (*P* = 0.0006; *r* = –0.31) with a more active AKT signature/pathway in EwS patients (**Supp. Fig. 4c**).

**Figure 4.**
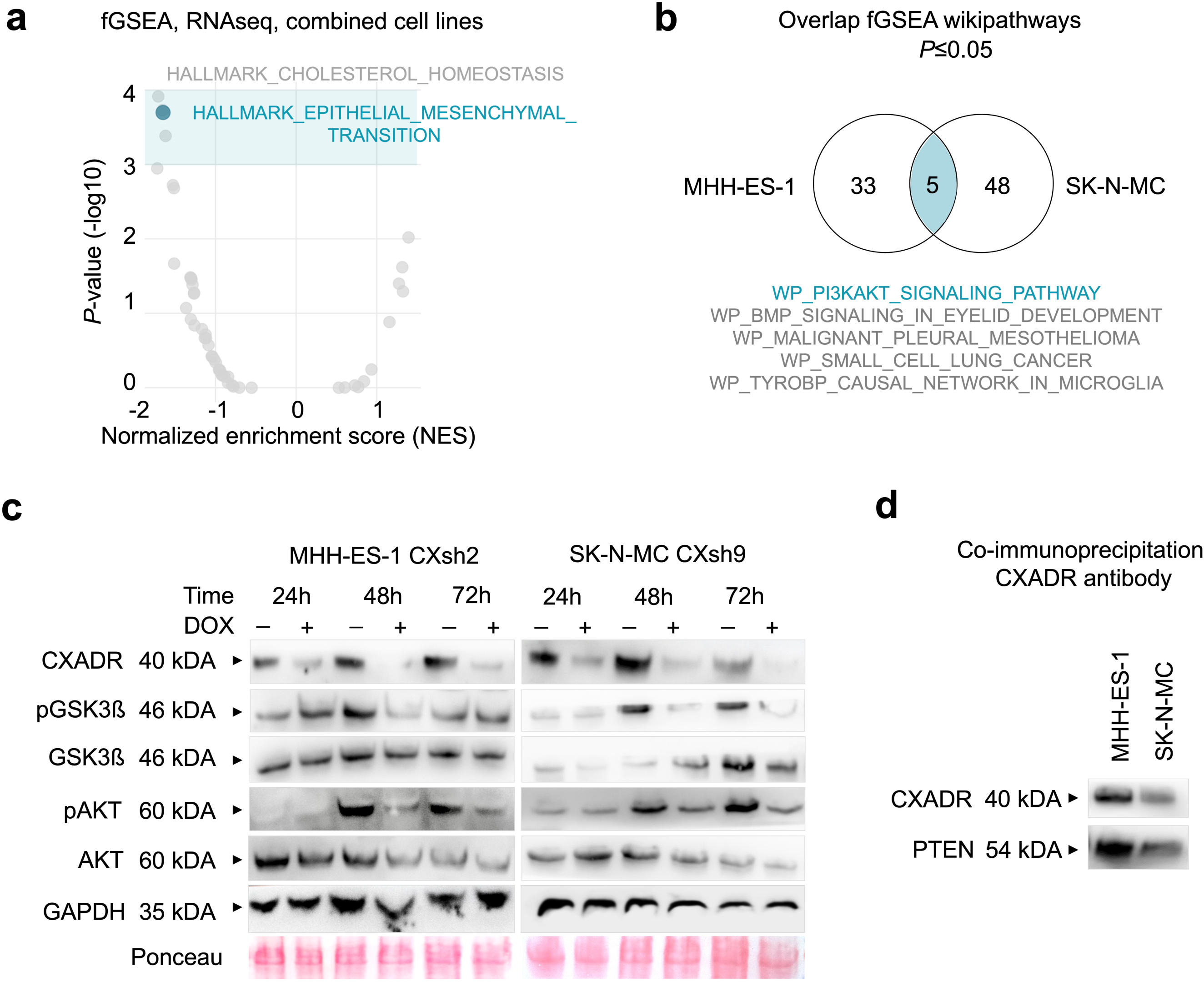
CXADR mediates EMT-phenotypes via the AKT signalling pathway. **(a)** fGSEA analysis on RNAseq samples from two EwS cell lines upon *CXADR* KD. Each dot represents a pathway from the Hallmark gene set. Blue area depicts pathways with *P* ≤ 0.001. Blue dot depicts the EMT hallmark signature. **(b)** fGSEA analysis on RNAseq samples from two EwS cell lines upon *CXADR* KD. Venn diagram depicts the significantly altered pathways in the Wikipathways gene set (C2 sub-set). **(c)** Immunoblot depicting time course experiments (24 h, 48 h and 72 h) for two EwS cell lines upon DOX-induced KD of CXADR. GAPDH and Ponceau were used as a loading controls. **(d)** Western blot analyses of MHH-ES-1 and SK-N-MC EwS cell lines co-immunoprecipitated using a CXADR antibody.

Overall, these findings highlight that CXADR forms a complex with PTEN and therefore participates in the regulation of AKT signaling, posing a therapeutic opportunity in EwS.

### Therapeutic targeting of AKT signaling synergizes with standard-of-care chemotherapy

To explore the translational potential of these findings, we employed a pan-AKT inhibitor (Capivasertib), which is clinically approved for the treatment of metastatic breast cancer in combination with Fulvestrant (Turner et al., 2023). We first carried out drug-response Resazurine assays and defined the IC_50_ of Capivasertib for four different EwS cell lines. Since we had previously shown that *CXADR* expression could be a prognostic marker for patient overall survival and that its high expression correlates with increased activity of the AKT pathway, we selected three cell lines (A-673, SK-ES-1, MHH-ES-1, and SK-N-MC, in ascending order regarding CXADR expression) that covered the full range of CXADR expression in EwS according to our ESCLA proteomic data (Orth et al., 2022) (**Fig. 5a**). Interestingly, IC_50_ analyses of Capivasertib showed that the level of CXADR expression was associated with the observed drug responses in all four tested cell lines, where higher CXADR expression resulted in better response (IC50_A-673_ = NA/no response; IC50_SK-ES-1_ = 5.2 µM; IC50_MHH-ES-1_ = 3.384 µM; IC50_SK-N-MC_ = 2.051 µM) (**Fig. 5b**). In line with these results, clonogenic assays using A-673, MHH-ES-1, and SK-N-MC showed that A-673 – the EwS cell line with lowest CXADR protein expression – did not respond to Capivasertib, even at relatively high doses (up to 12 µM) (**Fig. 5c**). Conversely, MHH-ES-1 and SK-N-MC, which exhibit a higher CXADR expression (**Fig. 5a**), showed a dose-dependent inhibition of clonogenic growth (**Fig. 5c**).

**Figure 5.**
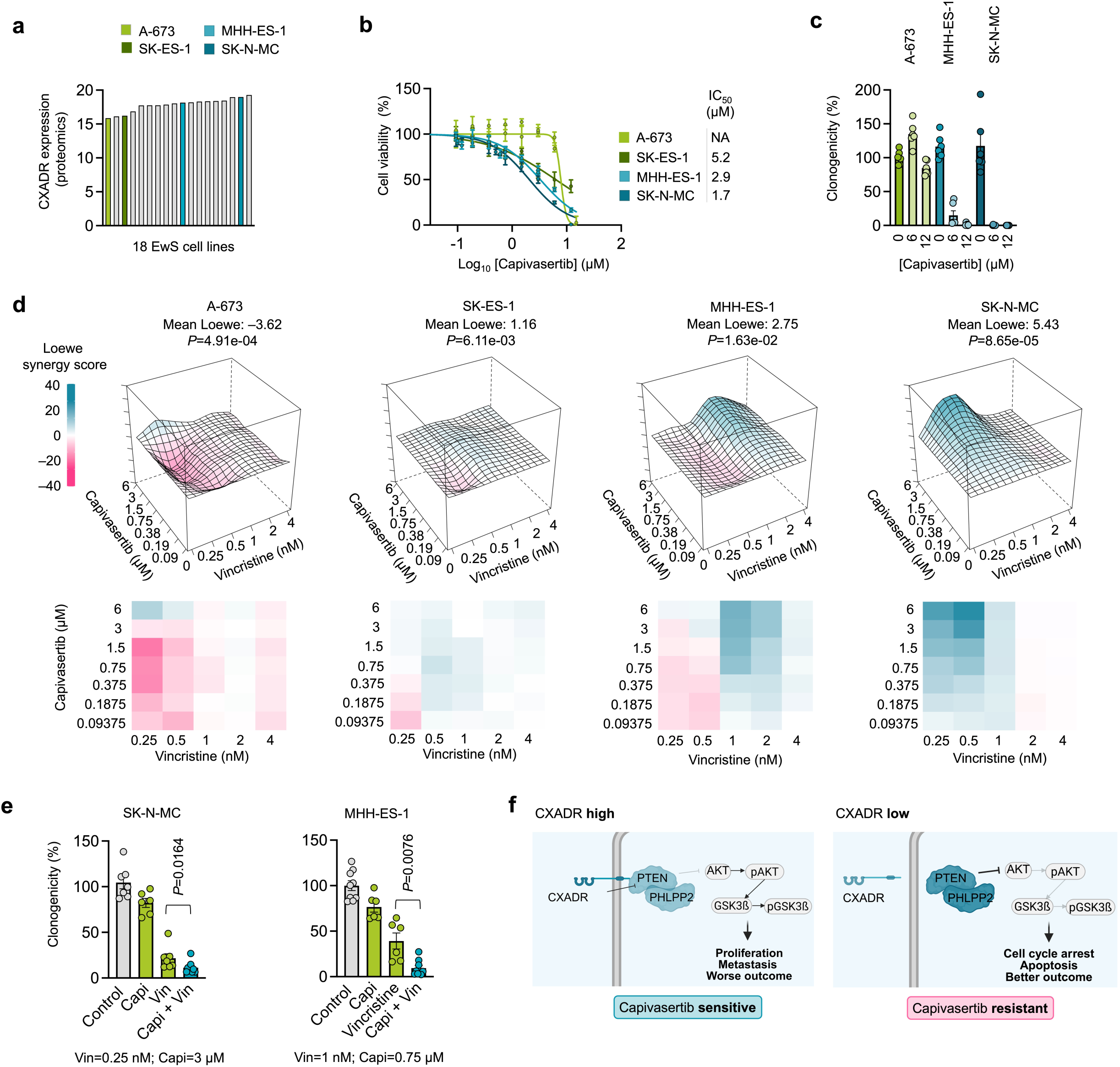
Therapeutic targeting of AKT signaling synergizes with standard-of-care chemotherapy. **(a)** CXADR protein expression in 18 EwS cell lines from the ESCLA dataset (Orth et al, 2022). **(b)** Resazurin cell viability analysis upon treatment with different concentrations of Capivasertib for 72 h in A-673, SK-ES-1, MHH-ES-1, and SK-N-MC cell lines. The calculated IC_50_ values are displayed on the right side. **(c)** Clonogenic growth assay using three EwS cell lines at different Capivasertib concentrations (0–12 µM). *n* = 6 independent biological replicates. Error bars represent SEM. **(d)** Loewe analysis of A-673, MHH-ES-1, and SK-N-MC treated with a combination of Vincristine and the AKT-inhibitor Capivasertib in vitro (blue indicates synergy). Summary level (mean) data from *n* = 4 biologically independent experiments. **(e)** Colony forming assays of MHH-ES-1 and SK-N-MC cell lines treated with a combination of Vincristine and Capivasertib. *n* ≥ 6 biologically independent experiments. Error bars represent SEM. Two-sided Mann-Whitney test. **(f)** Schematic illustrating key findings of this study.

Since Capivasertib is rarely used as a single agent, we tested it in combination with standard of care chemotherapy for EwS to evaluate a possible synergy effect. Doxorubicin, as well as Vincristine, were tested in combination with Capivasertib in different concentrations. While the combination with Doxorubicin showed little to no synergistic impact on all three cell lines tested (Mean Loewe_MHH-ES-1_ = 0.21, Mean Loewe_SK-N-MC_ = 1.46, Mean Loewe_A-673_ = –6.35) (**Supp. Figure 5a**), Vincristine showed a strong synergy with Capivasertib in SK-N-MC (Mean Loewe_SKNMC_ = 5.43), a moderate to high synergy in MHH-ES-1 (Mean Loewe_MHH-ES-1_ = 2.75), and no or lowly synergistic effect in A-673 (Mean Loewe_A-673_ = –3.62) and SK-ES-1 (Mean Loewe_SK-ES-1_ = 1.16), respectively, further supporting the concept that CXADR levels are a determinant for the efficacy of this combination therapy (**Fig. 5d**). To further test this phenotype in long-term assays, MHH-ES-1 and SK-N-MC cell lines were treated with Vincristine and Capivasertib as single agents as well as in combination for 10–12 days in clonogenic growth assays. Excitingly, these experiments demonstrated that the combination treatment achieved, on average, a 2–3-fold increase in efficacy in both cell lines as compared to the strongest single-agent effect (**Fig. 5e**).

Collectively, these results demonstrate that CXADR is a key regulator of the AKT pathway in EwS, and that AKT-inhibitors such as Capivasertib act synergistically with EwS standard-of-care chemotherapy in CXADR-high conditions. Our results support the concept that in EwS, CXADR may be employed as a biomarker to stratify patients who could benefit from including Capivasertib into their therapeutic regimen (**Fig. 5f**).

## DISCUSSION

Under the current treatment regimes, EwS patients with metastatic or recurrent disease have poor outcomes despite the application of highly toxic therapies (Setty et al., 2023). Increasing evidence supports the need for novel targeted therapies that may include personalized medicine approaches. Here, we show that patients displaying higher intratumoral CXADR expression at diagnosis have worse overall survival and an increased risk for metastasis, which is supported by our functional and translational in vitro and in vivo experiments.

Our finding that the tight-junction protein CXADR is involved in EMT-like processes is corroborated by its role as a positive regulator of the PI3K-AKT pathway. In fact, CXADR has also been linked to PI3K-AKT signaling in breast carcinoma, where canonical EMT processes are active (Nilchian et al., 2019). In EwS, preclinical inhibition of IGF-1R, an upstream regulator of PI3K-AKT signaling, in combination with standard-of-care chemotherapy led to a reduction of tumor growth, likely due to an increase in apoptosis (Benini et al., 2001; Martins et al., 2006; Scotlandi et al., 2005). Building on this knowledge, a phase II study enrolling 115 not otherwise preselected EwS patients with recurrent or refractory disease found that R1507, an anti-IGF-1R monoclonal antibody, displayed modest activity with an overall response rate of 10%. The drug was generally well tolerated, but the benefit appeared concentrated in a minority of patients (Pappo et al., 2011). Similarly, a recent randomized phase III trial including 298 eligible non-preselected patients found that adding the IGF-1R antibody ganitumab to interval-compressed chemotherapy for newly diagnosed metastatic EwS did not significantly improve event-free survival or overall survival compared with chemotherapy alone (DuBois et al., 2023). It should be noted that both studies acknowledged the need for identification of markers that are predictive of a benefit to fully capitalize on this approach. In line with this, a third phase I/II trial including 16 (phase I) and 107 (phase II) patients with advanced (relapsed/refractory) EwS evaluated the anti-IGF-1R monoclonal antibody figitumumab importantly including a pretreatment analysis of IGF-1 (total and free) in serum, and concluded that those patients with moderate to high concentration of circulating IGF-1 (2^nd^ and 3^rd^ quartiles, corresponding to 0.8–2.3 ng/mL free IGF-1) displayed improved overall survival upon anti-IGF-1R treatment (Juergens et al., 2011). These findings exemplarily demonstrate the importance of a pre-selection of patients to identify those who will benefit most from including a novel agent in their therapeutic regiment.

While direct targeting of PI3K-AKT in EwS remained unexplored, in recent years, several different compounds targeting this pathway have been developed, e.g. pan-PI3K inhibitors like Copanlisib (Munoz et al., 2021), pan-AKT inhibitors like Capivasertib (Shirley, 2024; Turner et al., 2023), or mTOR inhibitors like Everolimus (Jun et al., 2022; Raphael et al., 2020). Since single inhibition of individual PI3K-AKT components may lead to a compensatory upregulation of the signaling cascade or an interference with feedback loops within the pathway, PI3K-AKT inhibitors are often used in schemes that target multiple effectors of the signaling cascade or administered together with chemotherapy (Glaviano et al., 2023; Munoz et al., 2021; Shirley, 2024; Willis et al., 2024). In this study we used Capivasertib, a pan-AKT inhibitor, blocking all three variants of AKT, which has already been clinically approved by the FDA and EMA for treatment of metastatic breast cancer in combination with Fulvestrant, an estrogen-receptor antagonist (Shirley, 2024; Turner et al., 2023). It should be noted that while the A-673 cell line displayed the lowest expression of CXADR (**Fig. 5a**) which could explain the low Capivasertib efficacy, this cell line harbors an activating V600E BRAF mutation (Gouravan et al., 2018), that could account for at least part of the observed phenotype. To rule this possibility out, we included a second CXADR-low cell line (SK-ES-1), which similarly showed a reduced efficacy of Capivasertib, in concordance with its relative CXADR expression (**Fig. 5a,b**).

The observed CXADR-dependent efficacy of Capivasertib in EwS was additionally improved by its combination with standard-of-care chemotherapeutics, such as Vincristine, where again a dose-dependency on CXADR-expression was apparent. Although the precise mechanism for this synergy remains to be fully explored, it is tempting to speculate that the inhibitory effect of Vincristine on the mitotic spindle during mitosis may be potentiated by simultaneously blocking the PI3K-AKT signaling and thus further inhibiting tumor cell proliferation and promoting apoptosis.

Collectively, our results demonstrate that high CXADR can be used as a biomarker for patient stratification. Further, high CXADR sensitizes EwS cells for AKT inhibition, particularly in combination with EwS standard-of-care chemotherapy (**Fig. 5f**). In addition, this study presents a blueprint of how integration of in situ data with preclinical validation can help develop new potential strategies for personalized medicine.

## SUPPLEMENTARY FIGURE LEGENDS

**Supplementary Figure 1. Identification of CXADR as a potential key mediator for metastasis in EwS. (a)** Expression of *CXADR* in primary tumors versus metastases in the cohort of 166 EwS patients (second cohort). Two-sided Mann-Whitney test. **(b)** Overall survival exclusively displaying samples derived from primary tumors in a second cohort of 166 EwS patients according to *CXADR* expression. The best percentile was determined as the cutoff between groups. Log-rank (Mantel-Cox) test. **(c)** Overall survival was exclusively displayed for patients presenting with localized disease in a second cohort of 166 EwS patients, categorized by *CXADR* expression. The best percentile was determined as the cutoff between groups. Log-rank (Mantel-Cox) test.

**Supplementary Figure 2. CXADR drives tumorigenesis in EwS in vitro. (a)** siPOOL-mediated KD of *CXADR* in two EwS cell lines. *n* ≥ 4 independent biological replicates. *P* values were determined using a two-sided Mann-Whitney-test. **(b)** Proliferation assay after siPOOL-mediated KD of *CXADR*. *n* ≥ 4 independent biological replicates. Error bars represent SEM. Two-sided Mann-Whitney-test. **(c)** Remaining mRNA expression of *CXADR* after 72 h of DOX treatment using two additional different shRNAs in two EwS cell lines. n *=* 4 independent biological replicates. Error bars represent SEM. Two-sided Mann-Whitney test. **(d)** Cell cycle analysis of MHH-ES-1 and SK-N-MC cell lines with a shRNA-mediated conditional KD of *CXADR*. *n* = 3 independent biological replicates. Error bars represent SEM. *P*-values were determined using Student’s t-test.

**Supplementary Figure 3. CXADR drives primary tumor growth and metastatic spread in EwS in vivo. (a)** Ex vivo histological analysis of necrotic area of xenografted MHH-ES-1 and SK-N-MC cell lines upon *CXADR* KD. Two-sided Mann-Whitney test. **(b)** Kaplan-Meier plots of overall survival of animals orthotopically injected with MHH-ES-1 cells harboring a DOX-inducible *CXADR* KD. Log-rank (Mantel-Cox) test.

**Supplementary Figure 4. CXADR mediates EMT-phenotypes via the AKT signalling pathway. (a)** Bar plot of the 10 top hits following fGSEA analysis using the H gene set on RNAseq samples from two EwS cell lines upon *CXADR* KD. **(b)** Volcano plot depicts an independent fGSEA analysis using Wikipathways of MHH-ES-1 and SK-N-MC RNAseq upon *CXADR* KD. Blue area depicts pathways with *P* < 0.05, NES < –1. **(c)** Correlation analysis of the AKT_DOWN-signature in the second EwS patient cohort (*n* = 166). Simple linear regression model.

**Supplementary Figure 5. Therapeutic targeting of AKT signaling synergizes with standard-of-care chemotherapy. (a)** Loewe analysis of A-673, MHH-ES-1 and SK-N-MC treated with a combination of Doxorubicin and the AKT-inhibitor Capivasertib in vitro (blue indicates synergy). Summary level (mean) data from *n* = 3 biologically independent experiments.

## METHODS

### Provenience of cell lines and cell culture conditions

Human EwS cell lines MHH-ES-1 (RRID:CVCL_1411) and SK-N-MC (RRID: CVCL_0530) were provided by the German Collection of Microorganisms and Cell cultures (DSMZ). Human EwS cell line A-673 (RRID:CVCL_0080), human mesothelial cell line MeT-5A (RRID:CVCL_3749), human embryonal kidney cell line HEK293T (RRID:CVC_0063) and human fibroblast cell line MRC-5 (RRID:CVCL_0440) were purchased from the American Type Culture Collection (ATCC). All cell lines were grown under standard conditions at 37 °C and 5% CO_2_ in humidified atmosphere in RPMI 1640 medium or DMEM (MRC-5) with stable glutamine, supplemented with 10% tetracycline-free fetal calf serum (FCS), and penicillin at a concentration of 100 U/ml as well as streptomycin at a concentration of 100 µg/mL. Cells were tested by nested PCR for mycoplasma contamination and routinely SNP/STR-profiled.

### Generation of DOX-inducible shRNA constructs

Specific shRNAs targeting *CXADR* (MWG Eurofins Genomics or Sigma-Aldrich) or a non-targeting control were cloned in the pLKO-Tet-on-all-in-one system (Plasmid #21915, Addgene, Watertown, MA, USA). The corresponding oligonucleotide sequences are provided in **Supp. Table 2**. The backbone was digested using EcoRI and AgeI restriction enzymes (NEB, Ipswich, USA), and the respective oligonucleotides were ligated into the vector using a T_4_ ligase (NEB). Lentiviruses were produced in HEK293T cells using psPAX2 and pCMV-VSV-G as packaging plasmids. MHH-ES-1 and SK-N-MC EwS cells were infected with the respective lentiviruses and selected with 0.4 µg/mL (SK-N-MC) or 0.5 µg/mL (MHH-ES-1) Puromycin (Invivogen, San Diego, USA). Following the successful selection, single-cell clones were generated. KD efficacy of the respective clones was assessed by qRT-PCR 72 h after addition of DOX (1 µg/mL, Sigma-Aldrich), and only clones with a KD of < 20% remaining expression of *CXADR* were selected for functional assays. Details of the cloning protocol using the pLKO system have already been described previously (Musa et al., 2019; Orth et al., 2022). Primer sequences for cloning and RT-qPCR are provided in **Supp. Table 4**.

### Transient transfection of cell lines using small interfering RNAs (siRNA)

For transient transfection of cells to induce a siRNA-mediated knockdown, 0.7–2.5×10^5^ EwS cells (depending on the cell line) were seeded on a 6-well plate and transfected 24 h after seeding. The siPOOL against CXADR, as well as the negative control, were reconstituted in nuclease-free water to achieve a working concentration of 10 µM. On each well 400 µL transfection mix consisting of Lipofectamine RNAiMAX (Thermo Fisher Scientific, Waltham, USA), OptiMEM (Gibco, Waltham, USA), and the respective siPOOL or the negative control, were added dropwise to achieve a final siRNA concentration of 5 nM per well. Media was changed 24 h after transfection.

### Nucleotide extraction, reverse transcription, and quantitative real-time PCR (qRT-PCR)

Total RNA extraction was carried out utilizing the NucleoSpin® RNA kit (Macherey-Nagel, Germany). Next, 1 µg of RNA was reverse-transcribed using the High-Capacity cDNA Reverse Transcription kit (Applied Biosystems, USA). qRT-PCR reactions were performed using SYBR green Mastermix (Applied Biosystems, USA) mixed with diluted cDNA (1:10) and 0.5 µM forward and reverse primer (total reaction volume 15 µL) on a Bio-Rad Opus instrument (Bio-Rad, Hercules, USA) and analyzed using Bio-Rad CFX Manager 3.1 software. To calculate gene expression levels, the 2^−(ΔΔCt)^ method was used, normalizing the values relative to the housekeeping gene *RPLP0* as an internal control. Oligonucleotides required for the qRT-PCR experiments were purchased from MWG Eurofins Genomics as well as from Sigma-Aldrich and are listed in **Supp. Table 4**. The thermal conditions for qRT-PCR were as follows: initialization (95°C, 2 min) for one cycle; denaturation (95°C, 10 sec), annealing (60°C, 10 sec) and extension (60°C, 10 sec) for 49 cycles; denaturation (95°C, 30 sec), annealing (65°C, 30 sec) and extension (melting curve 65°C increasing 0.5°C every 5 sec until 95°C) for one cycle.

### Protein extraction and western blot

MHH-ES-1 and SK-N-MC cells containing shRNAs against *CXADR* were treated for 96 h with DOX (1 µg/mL) to induce the *CXADR* KD. Whole cellular protein was extracted by direct lysis with Laemmli sample buffer (pH 6.8, glycerin, 20% SDS, 1M Tris pH 7.5, bromphenol blue Na-Salt) (Applied Biosystems, USA), DTT (AppliChem, Germany), and Benzonase (Sigma-Aldrich, Germany). The extracted proteins were separated on a 10% SDS-PAGE gel at 100V and blotted on PVDF membranes. Antibodies are provided in **Supp. Table 5**. Protein detection was achieved using chemiluminescence and Immobilon Western HRP Substrate (Sigma-Aldrich, St. Louis, MO, USA).

### Proliferation assays

Proliferation assays were performed in analogy to Funk and Musa, 2021. Briefly, 1–1.5×10^5^ EwS cells containing either a DOX-inducible non-targeting control shRNA or *CXADR*-targeting specific shRNAs were treated with or without DOX (1 µg/mL) for 72 h and subsequently harvested and stained with Trypan blue (Sigma-Aldrich). Viable and non-viable cells were then quantified using a standardized hemocytometer (Merck KGaA, Darmstadt, Germany).

### Clonogenic growth assays

Depending on the cell line, 1–5×10^3^ cells containing either a DOX-inducible *CXADR*-specific or non-targeting control shRNA were treated with or without DOX (1 µg/mL) for 10–14 d. Then, cells were fixed with 100% methanol at –20 °C for 10 min and stained with crystal violet staining solution (2.3% CV solution, 4% formalin, 100% methanol, PBS) for 15 min. The number and area of the colonies were quantified on scanned plates using ImageJ (National Institutes of Health, MD, USA). The clonogenicity index was calculated by multiplying the number of colonies by the corresponding average colony area.

### 3D sphere assays

2.5×10^5^ EwS cells containing a DOX-inducible *CXADR*-targeting shRNA were plated per well on a low attachment cell-repellent 6-well-plate (Greiner Bio one, Frickenhausen, Germany) and treated either with or without DOX (1 µg/mL) for 14 d. The cells were then stained using 5mg/ml MTT (Sigma-Aldrich). Number and area of the spheres were quantified on scanned plates using ImageJ. The sphere index was calculated by multiplying the number of spheres by the corresponding average sphere area.

### Cell cycle analysis

Depending on the cell line, 1–1.5×10^5^ EwS cells per well containing either a DOX-inducible *CXADR*-targeting or non-targeting control shRNA were treated either with or without DOX (1 µg/mL) for 72 h. Then, cells were harvested, washed with cold PBS, and afterwards fixed by slowly adding ice-cold 70% ethanol to the cell pellet while vortexing. Cells were treated with 100 µg/mL RNAse (Applied Biosystems, USA) and stained with 50 µg/mL propidium iodide (PI) (Sigma-Aldrich). Samples were measured by BD FACSCanto II (BD Biosciences, Franklin Lakes, NJ, USA), and results were analyzed using FlowJo (BD Biosciences).

### Human samples and ethics approval

Human samples in the first cohort were retrieved and analyzed from the Cooperative Ewing Sarcoma Study (CESS) with approval of the ethics committee of the Medical Faculty of Heidelberg University (approval ID S-211/2021). All patients gave informed consent. Human samples in the second cohort correspond to public microarray data of 166 primary EwS tumors (GSE63157, GSE34620, GSE12102, GSE17618) and were analyzed as described in Musa et al., 2019.

### RNA sequencing (RNA-seq)

RNA quality was assessed by the Agilent 2100 Bioanalyzer before library preparation. After, 2 µg of high-quality RNA were prepared for sequencing. Library preparation was done according to the TruSeq RNA Exome (Illumina) protocol. The workflow included purification and fragmentation of mRNA, first and second strand cDNA synthesis, end repair, 3’ ends adenylation, ligation of adapters, PCR amplification, quantification of libraries, normalization, and library pooling. Between the mentioned steps, cleanup cycles were introduced. The sequencing was performed on a NovaSeq 6000 S1/S4 (Illumina). NovaSeq 6000 was performed with 100 bp ‘paired-end’ sequencing technology employing high-output flow cells. All samples were aligned to the genome using STAR aligner and counts were further processed using the DESeq2 package in R (version 1.46.0, https://bioconductor.org/packages/release/bioc/html/DESeq2.html).

### In vivo experiments

For evaluation of local tumor growth using subcutaneous xenografts in vivo, 2×10^6^ MHH-ES-1 or SK-N-MC EwS cells containing a specific shRNA against *CXADR* were resuspended in HBSS (Corning, Corning, NY, USA) and mixed with GeltrexTM (LDEV-Free, hESC-Qualified, Reduced Growth Factor Basement Membrane Matrix (Matrigel)) (Thermo Scientific, Waltham, MA, USA) in 1:1 proportion. After, the mixture was slowly injected subcutaneously in the right flank of 10–12 week-old NOD/scid/gamma (NSG) mice as previously described (Cidre-Aranaz and Ohmura, 2021). Tumor diameters were measured three times per week using a caliper, and tumor volume was calculated by the formula (L×I^2^)/2, where L is the length and I the width of the tumor. When the tumors reached an average volume of 50 mm^3^, the animals were randomized into two groups. The mice in the control group received 17.5 mg/mL sucrose (Carl Roth GmBh + Co KG, Karlsruhe, Germany) in drinking water. In contrast, the mice in the experimental group were treated with 20 mg/mL DOX (Thermo Fisher, Waltham, MA, USA) dissolved in drinking water with 50 mg/mL sucrose to induce an in vivo KD of *CXADR*. Once the experimental endpoint was reached, animals were sacrificed by cervical dislocation. General humane endpoints were defined as follows: body weight loss of 20%, apathy, piloerection, self-isolation, aggression as a sign of pain, self-mutilation, motor abnormalities such as a hunched back and reduced motor activity, as well as any other unphysiological or abnormal body posture, as well as breathing difficulties. To study the growth of EwS cell lines directly in the bone and their metastatic potential, EwS cells were orthotopically injected in the tibial plateau of NSG mice. Before the injection, the mice were treated with anesthesia and analgesia using inhaled isoflurane (1.5–2.5% in volume) and subcutaneous injection of Metamizole (800 mg/kg body weight). The eyes of the animals were protected with Bepanthen eye cream. Then, 2×10^5^ MHH-ES-1 and SK-N-MC EwS cells harboring an inducible shRNA-mediated KD of *CXADR* were injected into the right proximal tibial plateau of NSG mice dissolved in 20 µL HBSS using a 30 G needle (Hamilton, USA). For further pain prophylaxis following the injection, mice subsequently received Metamizole in their drinking water (800 mg/kg mouse weight) for 24 h. At the first day after injection, the mice were randomized in two groups. One group received 2 mg/mL DOX with 50 mg/mL sucrose dissolved in drinking water to induce the shRNA-mediated *CXADR*-KD whereas the other group only received sucrose in drinking water. All mice were closely monitored every two days and weighed once per week. Tumor growth was measured using a caliper. All mice were sacrificed by cervical dislocation before the primary tumors reached the maximum size, or if the animals exhibited the first signs of limping at the injected leg or met any of the humane or general endpoints as listed above. Afterwards, subcutaneously or orthotopically xenografted tumors and metastases were extracted. A section of each of the primaries or metastases was resected, snap frozen in liquid nitrogen and reserved for RNA or protein extraction to confirm the efficiency of the *CXADR* KD and the levels of the fusion. The remaining parts of the tumors and metastases were formalin-fixed paraffin-embedded (FFPE) for (immuno)histological analysis. All animal experiments were approved by the government of North Baden (permit number G3-20) and conducted under ARRIVE guidelines and recommendations of the European Community (86/609/EEC) and UKCCCR (guidelines for the welfare and use of animals in cancer research).

### Immunohistochemistry

Paraffin-embedded tissue sections (3–4 µM) were deparaffinized and rehydrated in distilled water. Antigen retrieval was performed using a microwave (CXADR, Cleaved Caspase 3 (CC3)) or a steamer (Ki67) with Citrate Buffer (pH 6.0) for 20 sec at 98 °C, followed by cooling down to room temperature (RT). After rinsing with Tris Buffered Saline with 0.05% Tween-20 (TBST, Sigma-Aldrich), sections were blocked with blocking solution BLOX ALL (Vector Laboratories, Newark, USA) for 15 min at RT. Subsequently, the sections were incubated with 2.5% (CXADR) or 2% (CC3, Ki67) horse serum (Vector Laboratories) for 25 min (CXADR) or 10 min (CC3, Ki67) at RT to minimize non-specific binding. After two washing steps with TBST, sections were incubated with the primary antibody against CXADR (rabbit anti-CXADR, concentration 1:25, abcam, #ab223689), CC3 (rabbit anti-Cleaved-Caspase 3, concentration 1:100, cell signaling, #9661) or Ki-67 (rabbit anti-human Ki-67, concentration 1:200, cell mark #275R-16, clone SP6) in DAKO antibody diluent (Agilent technologies, Santa Clara, USA) for 2 h at 37 °C. After three washing steps with TBST, sections were incubated with a secondary horseradish peroxidase (HRP)-coupled horse-anti-rabbit antibody (lmmPRESS, HRP Universal PLUS Polymer Kit) (Vector Laboratories) for 30 min at RT. After three washing steps with TBST, chromogen staining was performed using 3,3’-diaminobenzidine (DAB) for 10 min at RT. Then, sections were rinsed in distilled water for 5 min, followed by tap water for 10 min. Counterstaining was performed using Haemalaun (Carl Roth GmbH + Co. KG, Karlsruhe, Germany) for 1 min, followed by rinsing in tap water for 10 min. Finally, sections were dehydrated, cleared, and mounted with Aquatex (Merck KGaA).

HE stains of EwS xenografts were performed according to routine protocols. The slides were scanned and digitalized afterwards using the NanoZoomer-SQ Digital Slide Scanner (Hamamatsu Photonics K.K., Shizuoka, Japan) and visualized using NDP.view2 image viewing software (Hamamatsu Photonics K.K.). After, mitoses of EwS cells in the xenografts were quantified by a blinded observer in 5–10 high-power fields (HPF) per sample.

### Fast gene-set enrichment analysis (fGSEA) and single-sample GSEA (ssGSEA)

Using the cohort of 166 EwS patients, all genes were Pearson-correlated with the expression of *CXADR*. The entire list of genes with their respective correlation score was imported into R and analyzed using the fGSEA package in R (version 1.32.4, https://bioconductor.org/packages/release/bioc/html/fgsea.html) with different gene sets from MSigDB. For further analysis, the data from the RNA-seq were imported into R following the DESeq2 analysis. Again, gene sets from MSigDB were used and fGSEA was performed in R. Results were visualized using the ggplot2 package (version 3.5.2, https://ggplot2.tidyverse.org). To check individual GSEA scores on the cohort of 166 EwS patients, ssGSEA (single sample GSEA) was performed in R using the GSVA (version 2.0.7, https://bioconductor.org/packages/release/bioc/html/GSVA.html) and GSEABase (version 1.68.0, https://www.bioconductor.org/packages/release/bioc/html/GSEABase.html) package in R.

### Drug-response assays

For initial drug tests, EwS cells harboring a DOX-inducible shRNA-mediated KD against *CXADR*, wild-type EwS cell lines with different endogenous levels of *CXADR* or non-cancerous control cell lines (MRC-5, Met-5A) were seeded in a 96-well plate (1,000–5,000 cells per well, depending on the cell line). Cells containing shRNA were treated either with or without DOX for 24 h prior to drug treatment. One day after seeding, the respective drug was added to the plate. For each plate, an equal DMSO control corresponding to the highest concentration of DMSO used on the plate and cells without treatment were plated as well. After 72–96 h of treatment (depending on the experiment), 20 µL of resazurin (diluted 1:10) were added per well and incubated for 6–8 h (depending on the cell line). Readout was done on a plate reader. For clonogenic growth assays, 1,000–5,000 cells (depending on the cell line) were seeded in duplicates on a 6-well plate. 48 h after seeding, drug treatment was started with two different concentrations and one control condition in duplicates. The drug was refreshed every 48 h. Plates were stained 10–14 days after seeding using crystal violet.

### Synergy assays

To access possible synergistic effects between drugs, wild-type EwS cell lines with different endogenous levels of *CXADR* were seeded on a 96-well plate (1,000–5,000 cells per well, depending on the cell line). 24 h after seeding, the two drugs were added to the plate. 72–96 h after the beginning of treatment, resazurin was added to the plates and readout was done using a plate reader. Synergistic effects were determined using the SynergyFinder package (version 2.2.0, https://www.bioconductor.org/packages/release/bioc/html/synergyfinder.html) in R. Synergetic effects were calculated using the Loewe synergy model.

For colony growth assays using a drug combination, 1,000–5,000 cells (depending on the cell line) were seeded per well on a 6-well plate. Drugs were refreshed every 48 h, and plates were stained 10–14 d after seeding using crystal violet.

### Co-Immunoprecipitation (Co-IP)

Per Co-IP 1.5 mg Dynabeads (Thermo Scientific, Waltham, MA, USA) were coupled with 10 µg *CXADR* antibody at 37 °C overnight. Beads were washed according to the manufacturers’ protocol. For cell lysis, the buffer was prepared according to the manufacturers’ protocol and optimized using 100 mM NaCl, 2 mM MgCl_2_, and EDTA-free protease inhibitor. Cells were harvested and washed with PBS. The cell pellet was weighed, and a weight-adopted amount of lysate buffer was added to the pellet. The lysate was incubated with the *CXADR*-coupled Dynabeads for 30 min at 4 °C and washed multiple times. The purified protein complex was mixed with 6X reducing Laemmli buffer and loaded on a 10% SDS-PAGE gel. Western blot was performed as described above.

### Statistical analyses and software

Statistical data analysis was performed using GraphPad PRISM 10 or with R on the raw data. If not otherwise specified in the respective figure legend, comparison of two groups in in vitro experiments was carried out using a two-tailed Mann-Whitney test. If not otherwise specified in the figure legends, data are presented as dot plots with horizontal bars representing means and whiskers representing the standard error of the mean (SEM). The chosen sample size for all in vitro experiments was empirically selected with at least three individual and independent biological replicates. In Kaplan-Meier overall survival analyses, curves were calculated from all individual survival statistics of patients. The two groups were compared by the Mantel-Haenszel test to detect significant differences between them. Pearson’s correlation coefficients were calculated using Microsoft Excel. For the in vivo experiments, the sample size was determined using power calculations with ß = 0.8 and α = 0.05 based on preliminary data and compliant with the 3R system (replacement, reduction, refinement). Kaplan-Meier analyses of event-free survival were carried out by using GraphPad PRISM 10.

## Data availability

The data generated in this study are available in the German Human Genome-Phenome Archive (GHGA, data.ghga.de) under the GHGA accession number GHGAS43962012454044.

## ACKNOWLEDGEMENTS

We would like to thank Felina Zahnow, Stefanie Kutschmann, Sabrina Knoth, and Nadine Gmelin for excellent technical assistance. We are also grateful to Claudia Schmidt from the Light Microscopy Facility (LMF, German Cancer Research Center (DKFZ), Heidelberg, Germany) for her impeccable work in conducting immunohistochemical stains. We would also like to thank Gaby Blaser and Gabriele Schmidt from the LMF for additional support. We thank the DKFZ NGS Core Facility for providing excellent RNA-sequencing services.

## AUTHOR CONTRIBUTIONS

A.R., F.C.A., and T.G.P.G. conceived the study. A.R., F.C.A., and T.G.P.G. wrote the paper and drafted all figures and tables. A.R., M.Z., F.H.G., K.M.H., A.J., T.F., L.R.P., and P.T. performed in-vitro experiments and assisted with sample analyses. A.R. and M.Z. performed all bioinformatic and statistical analyses. A.R., M.Z., M.J.C.G., D.O. and F.C.A. carried out in-vivo experiments. F.C.A., R.I. and A.B. coordinated in-vivo experiments. U. D., E.A. and W.H. provided patient samples and/or clinical information. A.R. and T.G.P.G. performed immunohistological evaluation. J.S. provided guidance for bioinformatic analyses. F.C.A. and T.G.P.G. supervised the study and data analysis. All authors read and approved the final manuscript.

