## Supplementary figures and images for "Targeting CXADR-mediated AKT signaling suppresses tumorigenesis and enhances chemotherapy efficacy in Ewing sarcoma"

### Supp Figure 1

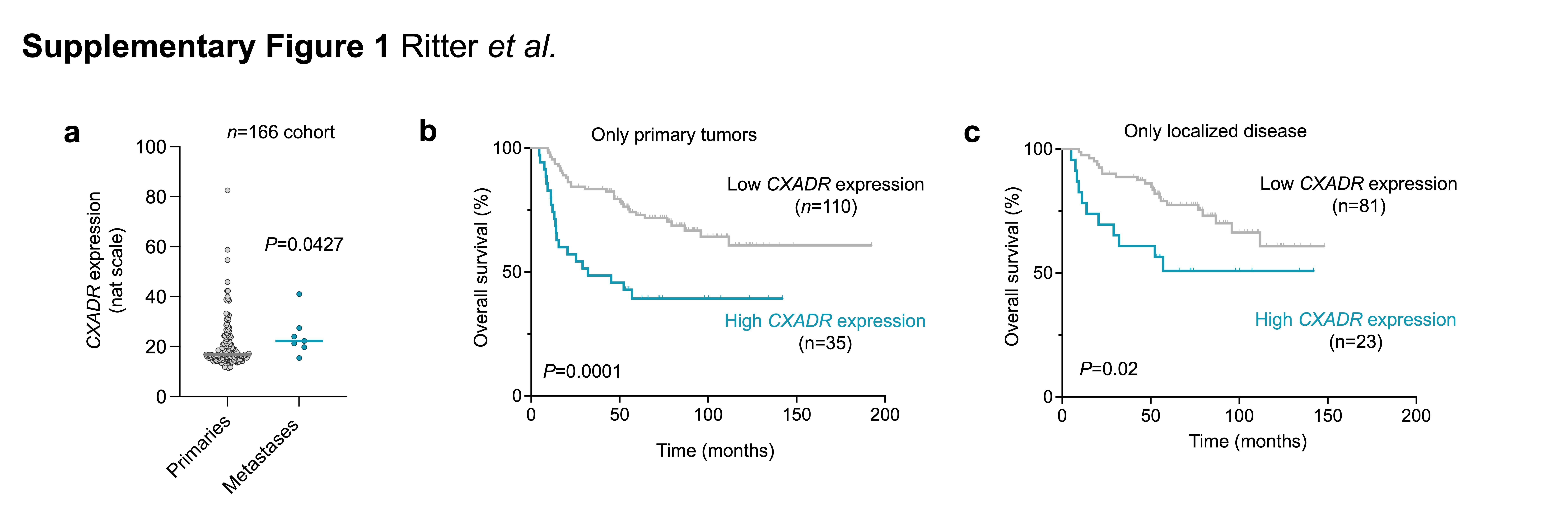

### Supp Figure 2

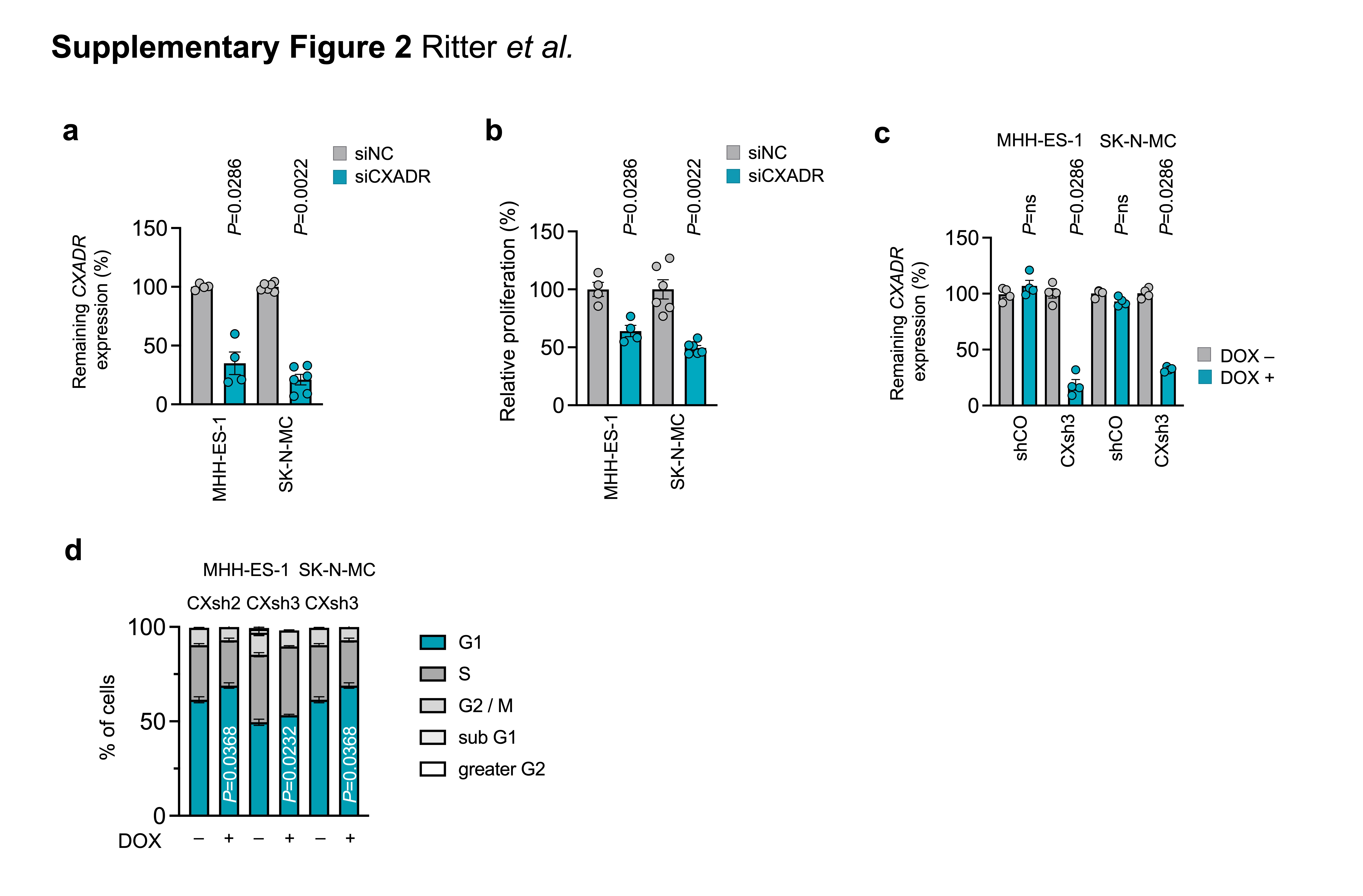

### Supp Figure 3

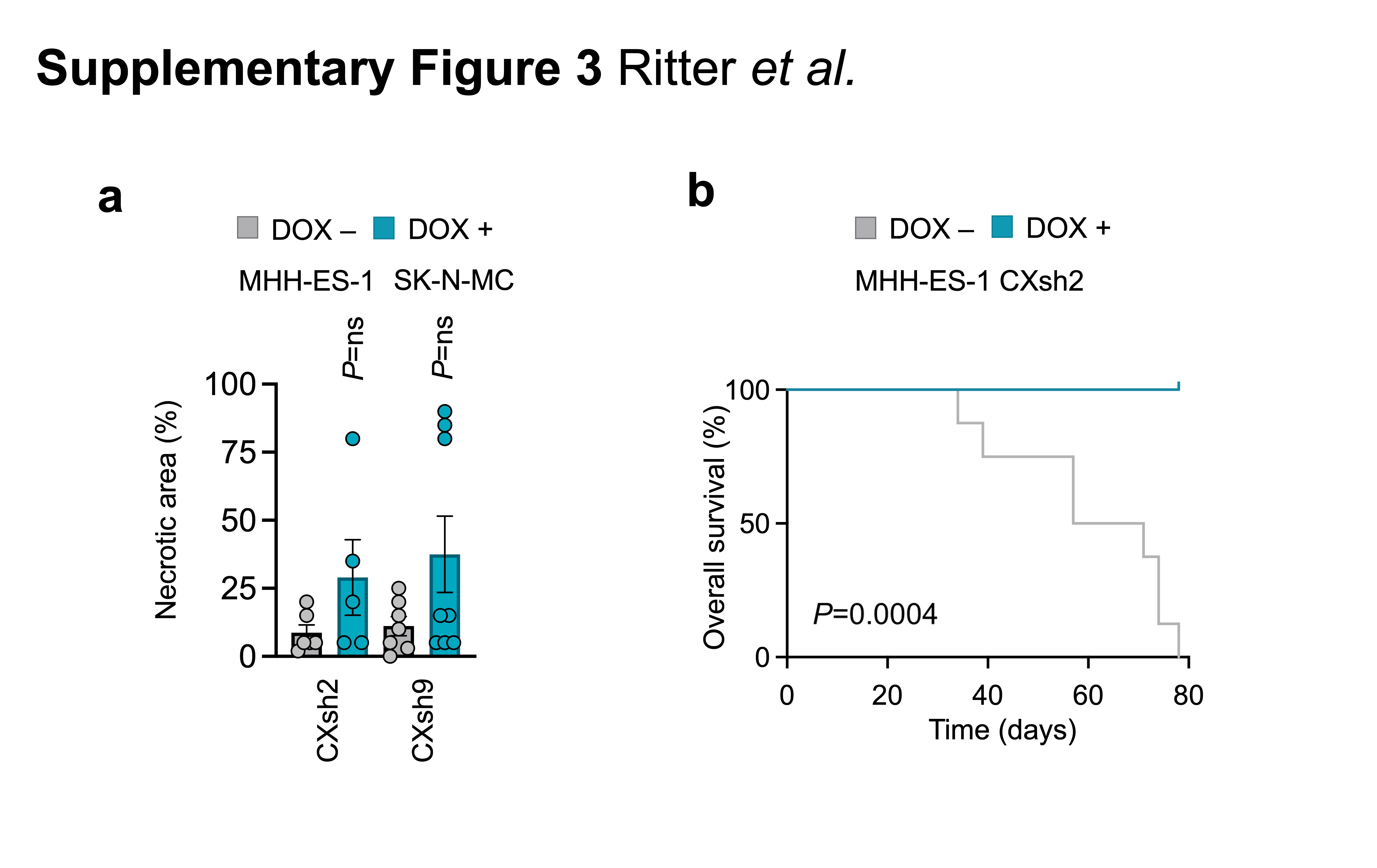

### Supp Figure 4

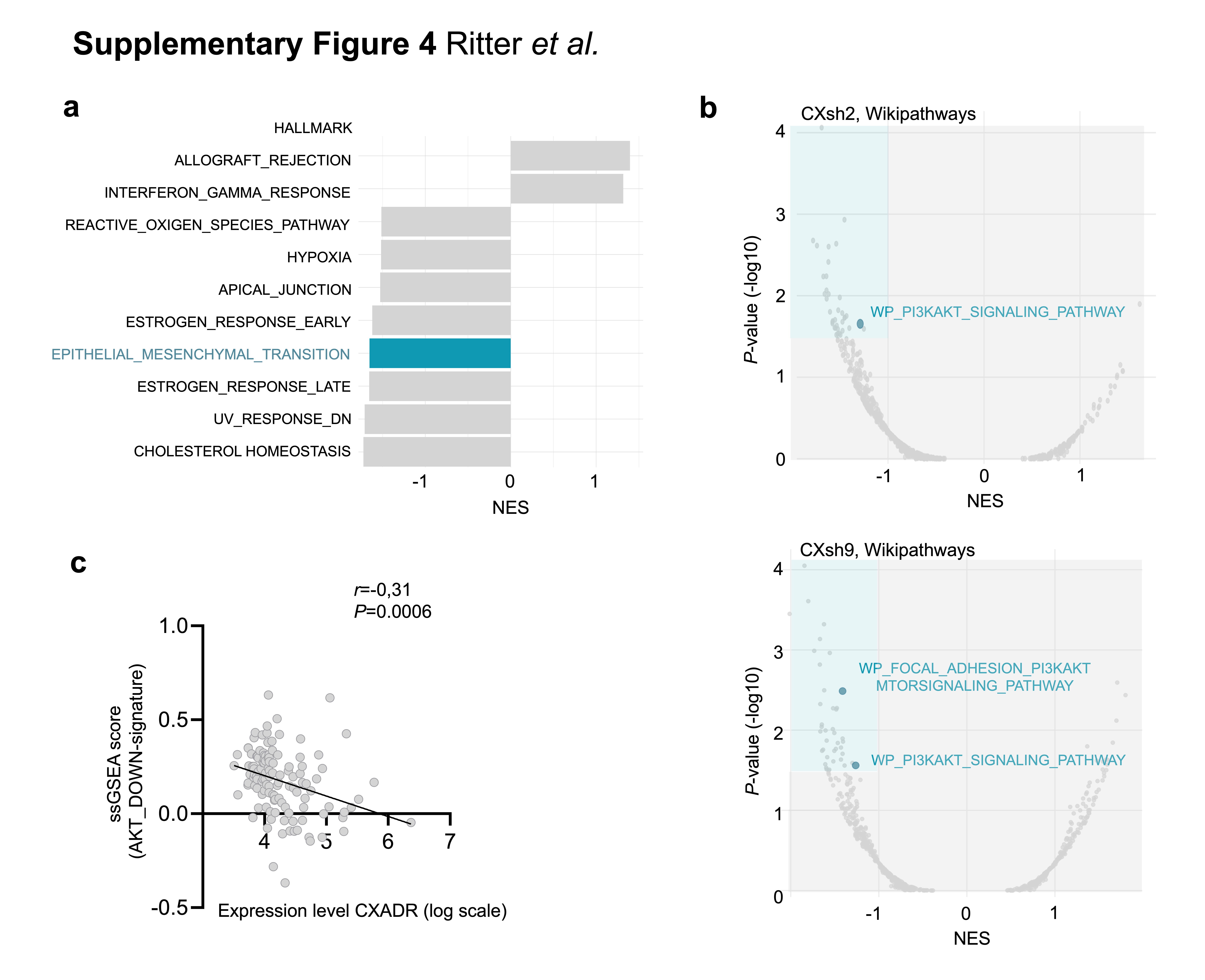

### Supp Figure 5

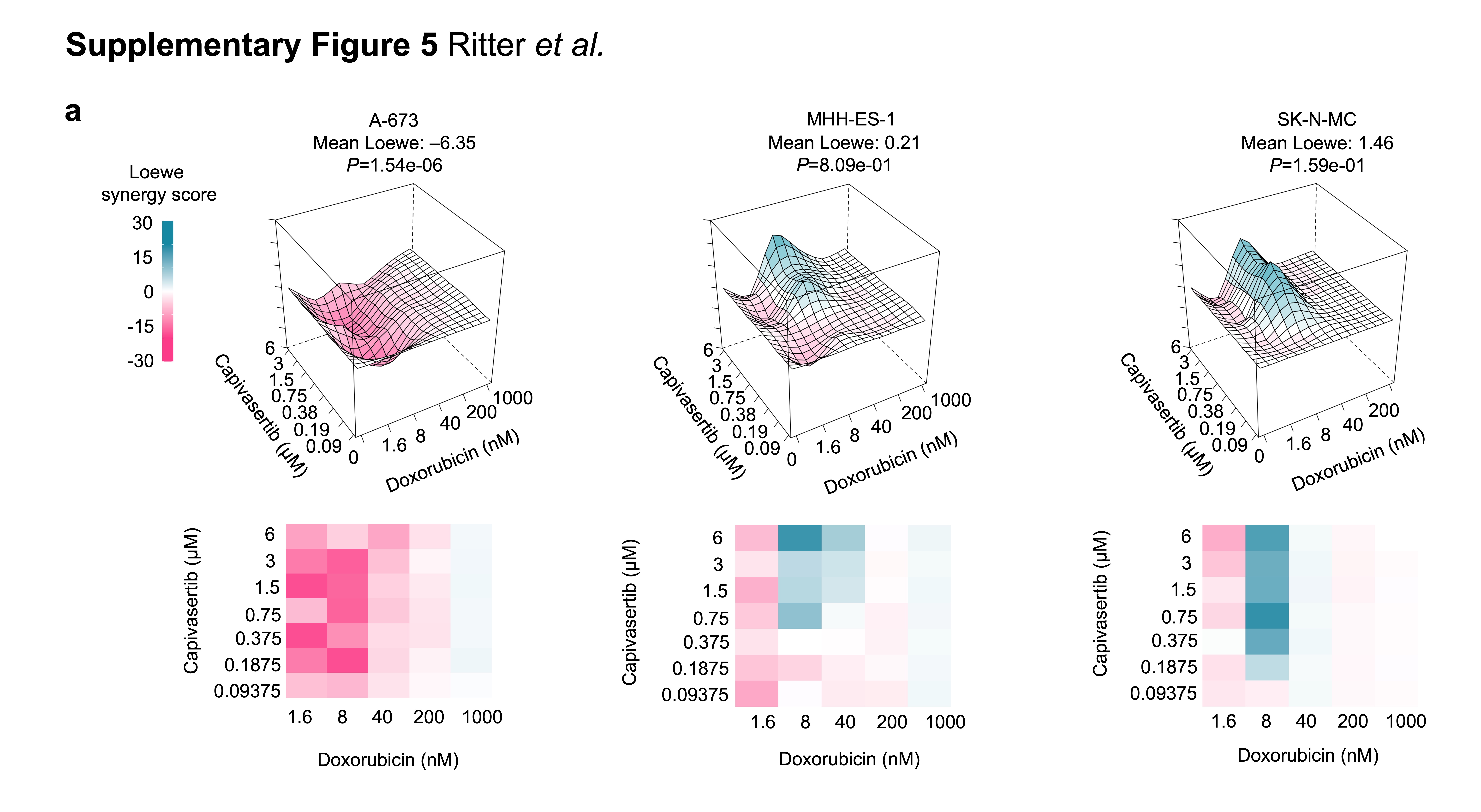
